# Network-based integration of gene expression and DNA methylation identifies prognostic biomarkers for early-stage pancreatic cancer

**DOI:** 10.64898/2026.02.09.704985

**Authors:** T Dhanushkumar, Krishnan Anbarasu, Sripad Rama Hebbar, Karthick Vasudevan, Karunakaran Rohini

**Affiliations:** Department of Biotechnology, School of Applied Sciences, REVA University, Bengaluru-560064, India; Department of Computational Biology, Saveetha School of Engineering, Saveetha Institute of Medical and Technical Sciences (SIMATS), Saveetha University, Chennai, Tamil Nadu, 602105, India; Manipal Academy of Higher Education (MAHE), Manipal, 576104, India; Institute of Bioinformatics, International Technology Park, Bangalore, 560066, India; Unit of Biochemistry and Medical Education, Faculty of Medicine, AIMST University, Semeling, Bedong, Malaysia

**Keywords:** Pancreatic cancer, early-stage biomarkers, multi-omics integration, partial correlation networks, WGCNA, machine learning

## Abstract

Pancreatic ductal adenocarcinoma remains one of the most lethal malignancies, largely due to the absence of reliable early-stage biomarkers. Here, we present a network-based multi-omics framework that integrates gene expression and DNA methylation data through partial correlation analysis to uncover prognostic markers. Four distinct networks were constructed: gene expression co-expression, methylation-only, multiplex (inter-layer connections linking the same genes across omics layers), and monoplex (fused multi-omics). Weighted gene co-expression network analysis (WGCNA) was applied to each network to select non-redundant, topologically representative hub genes as features for machine learning classification. Models trained on cross-layer (multiplex) features achieved an ROC of 82%, compared with 50–60% using single-omics features alone. The most strongly associated genes with poor prognosis include TFCP2L1, DHX32, and NCK1.

## 1. Introduction

Pancreatic Cancer (PC) has been ranked as the 12^th^ most common cancer type globally, with the death rates ranking at the 6^th^ position due to the failure in diagnosing in early stages. Overall survival rates for pancreatic cancers ranged from 7% to 13% (Leiphrakpam et al., 2025). The main reason for the low survival rates is its failure to detect the cancer in early stages due to the nonspecific early symptoms, which often lead to diagnosis only after significant tumor progression in later stages with metastasis condition. However, if early signs of the PC were clinically detectable, the survival rates would have been higher (Singhi et al., 2019). Currently, CA19-9 is the only FDA-approved serum biomarker for PC (Liang & Tong, 2023).

CA19-9, also known as the sialyl Lewis A, is a carbohydrate (tetrasaccharide) molecule normally present on the tumor antigens like MUC1, MUC5AC, MUC16, and CD44 (Barkeer et al., 2018). This antigen at a cutoff value of 37 U/mL has a sensitivity of around 71–81% with 82–90% specificity (Yang et al., 2022). However, a major limitation of this marker is that it can only be detected in the population with the FUT3 gene, PC patients lacking this gene cannot be detected through CA19-9. Additionally, these biomarkers are also seen in pancreatitis, liver diseases, lung fibrosis, and also in cancers of colon, lung, or liver, reducing their specificity (Ballehaninna & Chamberlain, 2012). Due to this major limitation, there is a great need for a biomarker that is highly specific to the PC and can be detected in early stages, and the integration of genomics with artificial intelligence offers a powerful strategy to address this challenge.

Epigenetic modification, particularly DNA methylation, plays a crucial role in regulating the transcriptional activity of genes without altering the underlying DNA sequence, and it also influences tumor prognosis. Similarly, transcriptomics has been used in analyzing the prognostic patterns of many cancers. In this context, both epigenomic and transcriptomic data have been consistently applied to screen numerous cancer-associated hub genes, and the same strategy can be extended to identify prognostic biomarkers in pancreatic cancer in early condetions (Xu et al., 2019).

Combining these datasets into a single multi-omics framework can provide more accurate and comprehensive biological insights. Combining the two types of data into a single multi-omics dataset that stores both methylation and expression data is very challenging, and networking is one of the widely used approaches for merging the multiple omics data’s in to single data (Fraunhoffer et al., 2022).

Networking and mathematical modelling are the key strategies that are used in drawing connections between the gene-gene correlations, and these gene-gene networks can be drawn between the genes from expression data and methylation data in the form of graphs (Jiang et al., 2025). Applications of the Network-based analysis have been applied to many biological data in both descriptive and predictive, like calculating the scores between the genes, how a particular gene influences the expression of other genes, and predicting the gene signatures (Ko & Brandizzi, 2020). One of the important applications of this networking in artificial intelligence is predicting the highly correlated features for the training, which can be done through weighted gene co-expression analysis, where researchers can calculate highly co-expressing genes. And the features derived from this WGCNA will store unique information in each feature, and this strategy can yield an high accuracy, since all the features store unique information (Chai et al., 2021).

In this study, our objective is to combine both methylation and expression data into a multi-cross-layer dataset that stores both expression and methylation profiles and predicts the early-stage biomarkers for PC using machine learning models, trained from the features derived from the WGCN. A detailed workflow adapted for this study is illustrated as a flowchart in **Figure1**.

**Figure 1.**
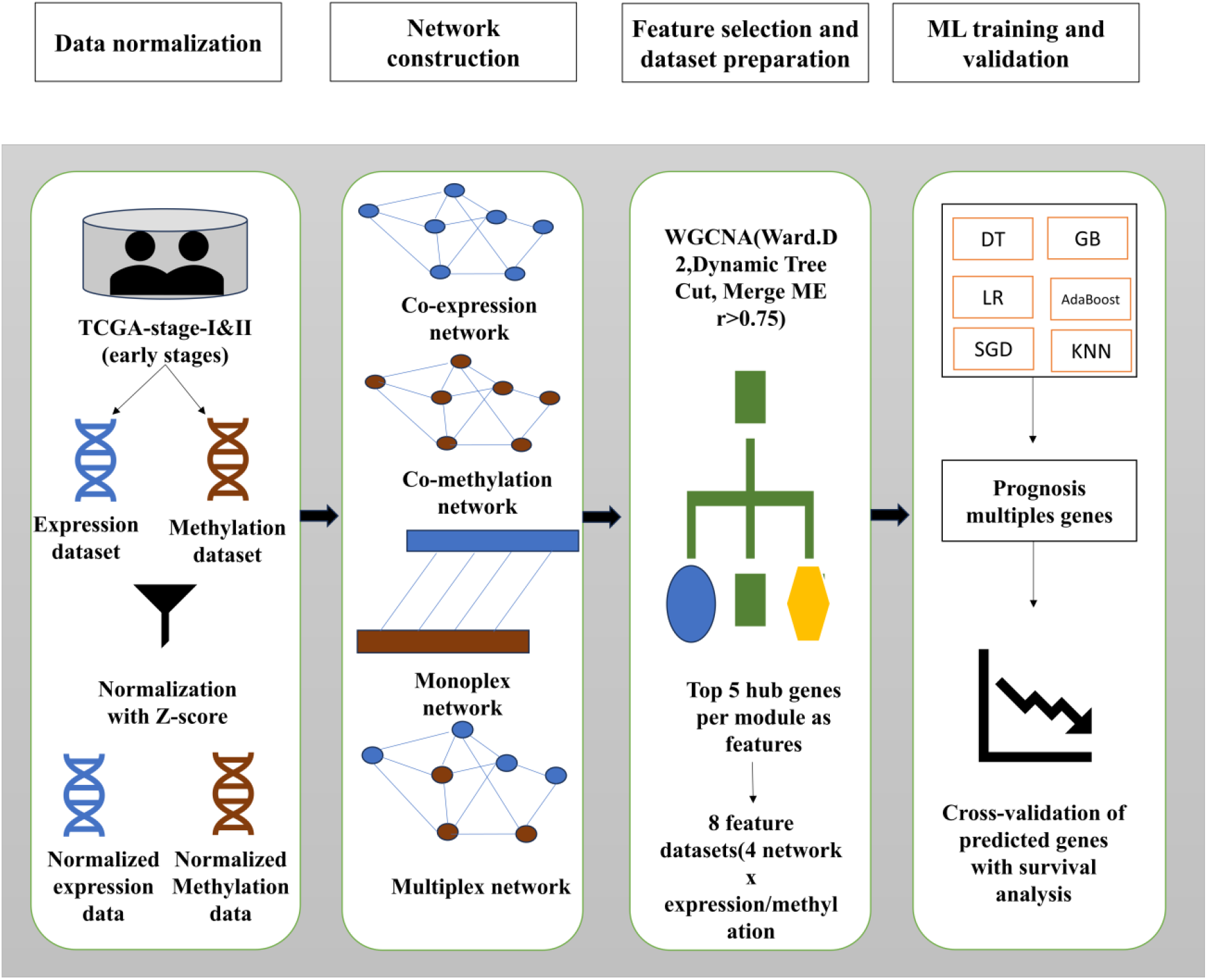
Ovewview of workflow adapted for this study

## 2. Methodology and methods

### Data acquisition and preprocessing

Transcriptomics and methylation omics data were gathered from the TCGA database. In TCGA, a total of 639 Transcriptomics samples and 528 DNA methylation samples were reported. Since the focus of our study is on early-stage patients, we selected 156 early-stage patients (1-a&b, 2-a &b) based on their clinical information and samples that shared common DNA methylation and expression data (Kim et al., 2024).

156 transcriptomics and methylation samples omics data were downloaded through gdc-clint tools, and the transcriptomics data were built using FPKM expression values (Jensen et al., 2017). Similarly, methylation data was downloaded through the gdc-clint tool, and the prob ID were matched to genes through IlluminaHumanMethylationEPICanno.ilm10b4.hg19. Each prob was mapped to its UCSC reference gene. Probes without valid gene annotations are discarded. After preparing the transcriptomics and methylation data, common genes between these two omics data were considered for network construction.

### Normalization and Network Construction

Normalization of the expression and methylation omics data was done using log10 and z-score methods. For the raw FPKM expression counts, First, a log10 transformation was applied to the raw expression values to reduce extreme variability and stabilize the distribution, since gene expression data typically contain highly skewed values with large dynamic ranges. Next, z-score normalization was applied to both expression and methylation datasets to scale all genes to a comparable range. Following normalization, genes with very little variation, which carry little information, were dropped from the analysis, and genes with high variability were considered (Cheadle et al., 2003).

A total of 4 different types of networks were constructed. To construct a network, genes were considered as nodes and the values as edges, and the network was constructed using the partial correlation method. The correlation will calculate the relation between the two genes, considering the effect of other genes (Kuijpers et al., 2021). The constructed 4 types of networks include:

- **Co-expression network**: In this network, gene–gene connections were derived exclusively from the expression matrix, meaning all nodes belong to the same molecular layer and the edges reflect expression-based associations only. Because the network is built from a single omics modality without integrating additional biological layers, it is characterized as a single-layer (single-omic) network.
- **Co-methylation network**: Similar approach as expression, a co-methylation network was constructed using only methylation values, where gene–gene connections were defined based on their methylation profiles. Because all nodes and edges originate from a single omics modality, this structure also represents a single-layer (single-omic) network.
- **Multiplex matrix**: In a multiplex network, each layer corresponds to one omics data type (gene expression or DNA methylation). The adjacency matrix of this multiplex network encodes the correlations within and between these layers. Specifically, intra-layer edges represent partial correlation between genes within the same data type (expression-expression or methylation-methylation), capturing direct gene-gene associations. The coupling between layers is represented by inter-layer edges connecting the same gene across expression and methylation layers, constructed using cross-layer partial correlations.
- **Monoplex matrix**: A Monoplex network was built by integrating both omics layers into a single fused network. Here, the adjacency matrix is constructed by combining the gene features across expression and methylation using the outer product between node properties. However, because the fusion results in one consolidated adjacency matrix rather than multiple interacting layers, the monoplex representation still functions as a single-layer network with mixed multi-omics information embedded within it.

### WGCNA analysis

WGCNA analysis was performed to screen the highly significant correlating features to train the machine learning (ML) models. Input for the WGCNA is the adjacent networks constructed in the previous step.

WGCNA uses hierarchical clustering to group genes into preliminary branches before module cutting. For this Ward.D2 method was used, which minimizes the total within-cluster variance during the clustering process. Ward.D2 works by merging clusters that result in the smallest increase in clustering error, making it more robust for gene expression data. It produces well-separated, compact clusters that serve as an excellent starting point for dynamic module detection. Post hierarchical clustering Module Identification was done Using Dynamic Tree Cut, where different modules with genes in similar functions were grouped, minimum genes in each module was set to 20. After generating the module, the modules with similar functions were merged into to single module using the module eigengene (ME). Modules with eigengene correlation above a threshold of >0.75 were merged (Langfelder & Horvath, 2007).

### Selection of highly significant features

To identify the most informative features, we selected highly significant hub genes from each WGCNA module based on two criteria: module membership (kME), which measures the correlation between each gene and the module eigengene, and intramodular connectivity, which indicates how strongly a gene is connected to other genes within the same module. Genes with both high kME and high connectivity were considered hub genes because they serve as central functional regulators that drive the biological behavior of each module. For model training, we built the dataset by choosing the top five hub genes from each module as predictive features. To achieve a complete representation of gene regulation, we included both gene expression and methylation data (Kosvyra et al., 2024).

A total of eight datasets were prepared from four networks: for each network, one dataset contained features based on expression data and another based on methylation data. This method significantly reduces the number of features, prevents confusion in module representation, and maintains strong biological relevance in subsequent machine learning analyses.

### Machine learning model training and evaluation

We have trained the prepared 8 different datasets on the 6 ML models, namely DecisionTree, GradientBoosting, LogisticRegression, AdaBoost, Stochastic Gradient Descent (SGD), and K-Nearest Neighbors (KNN). For the selected models, hyperparametric tuning was performed using the randomized search with cross-validation to increase the model’s accuracy and to prevent he models from overfitting. The training was made with an 80%-20% training and testing split.

1. For the Decision Tree classifier, hyperparameter optimization was performed across a search space of maximum depth values ranging from 2 to 200, and minimum sample requirements for node splitting ranging from 2 to 100. Tree splitting criteria were evaluated using *gini, entropy*, and *log_loss* measures with both *best* and *random* split strategies. Additionally, the proportion of features considered at each split was varied using options such as *log2, sqrt*, and continuous fractions between 0.3 and 1.0 (Gomes Mantovani et al., 2024).
2. For the Gradient Boosting classifier, tuning was conducted using the number of estimators ranging from 10 to 2000 and learning rates between 10^−4^ and 1. Tree depth was explored from 2 to 50, with subsampling ratios varied between 0.4 and 1.0. Minimum sample splits ranged from 2 to 50, and minimum samples per leaf from 1 to 48. The max-features parameter was optimized using options auto, sqrt, log2, and proportional values from 0.2 to 1.0 (Boldini et al., 2023).
3. For Logistic Regression, model optimization was performed over a regularization strength (C) range from 10^−5^ to 10^4^. Several solvers were compared, including lbfgs, liblinear, saga, newton-cg, and sag. Regularization penalties were tuned across *none, l1, l2*, and *elasticnet* options, with elastic-net mixing ratios varied from 0 to 1. Additionally, both unweighted and class-balanced configurations were evaluated (Zhang et al., 2018).
4. For the AdaBoost classifier, hyperparameter tuning was conducted using the number of estimators ranging from 10 to 1500, and learning rates between 10−410−4 and 10. Both SAMME and SAMME.R boosting algorithms were evaluated during tuning (Black et al., 2024).
5. The SGD classifier was optimized across multiple loss functions with regularization strengths (αα) ranging from 10−810−8 to 2, and iteration limits set between 500 and 20,000. Class weights were kept in balanced mode, and elastic-net mixing ratios varied from 0.01 to 1(Pan et al., 2023).
6. For K-Nearest Neighbors (KNN), optimization involved tuning the number of neighbors from 1 to 201, testing weight functions of uniform and distance. The power parameter pp for the Minkowski distance metric was varied from 1 to 5. Additionally, a variety of distance metrics were evaluated, including Minkowski, Euclidean, Manhattan, Chebyshev, Hamming, Canberra, and Bray-Curtis (Siddalingappa & Kanagaraj, 2023).

The accuracy and ROC curves for each dataset were created for comparison. Datasets with the highest accuracy were used to identify biomarkers associated with poor patient survival.

## 3. Results

### Data preprocessing and

Log10-transformed expression values ranged from 0 to 23.89285, and the raw methylation data values ranged from 0.003051967 to 0.9795942. Since the data values were not in the same range, the z-score was applied to both datasets. After applying the z-score, the expression data ranged from −6.764049 to 11.02749, and methylation ranged from −9.476452 to 12.26373. Therefore, normalization brought both omics datasets into a comparable scale, ensuring equal contribution during downstream analysis. Variance distribution plots of before and after normalization are illustrated in **Figure 2**.

**Figure 2.**
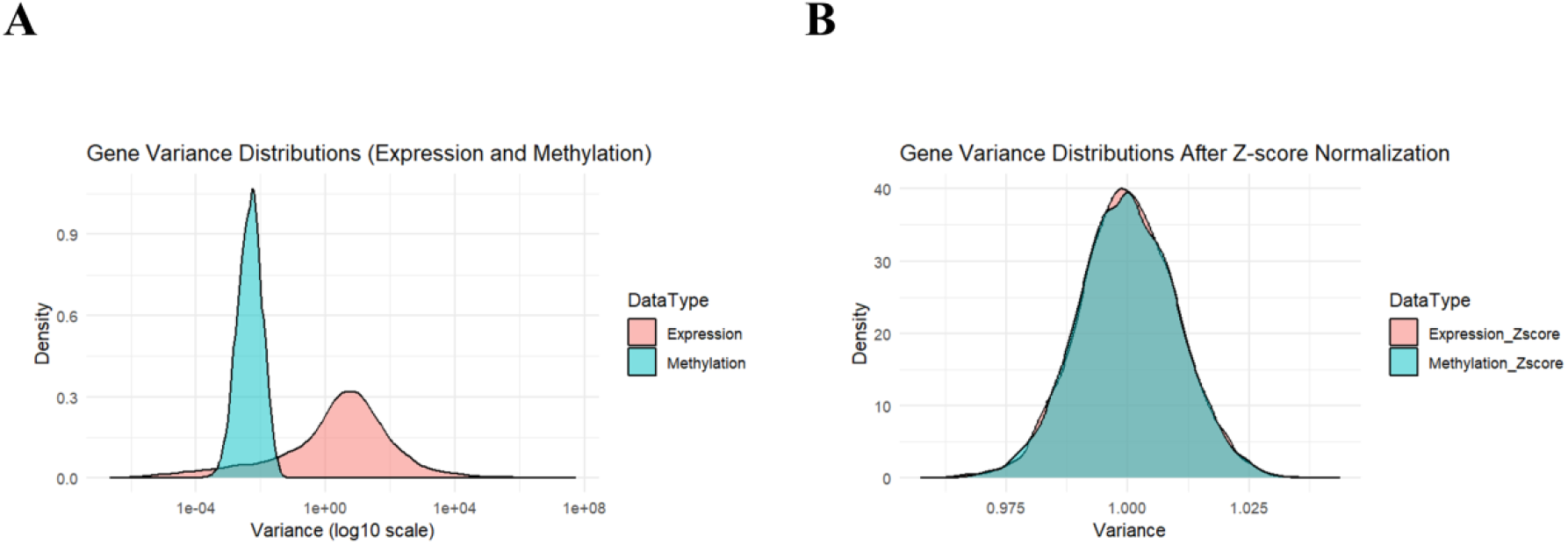
Before and after normalization of expression and methylation data using Log10 and the Z-score method

### Network construction

Using the normalized data, the top variance was selected using the rowVars function from the matrixStats package, ensuring that noise was reduced and only informative genes were retained. After selecting high variant genes, only 5707 common genes were found shared between both datasets, and these genes were used to construct the network. Since we have used genes as nodes and values as edges, the total number of nodes is 5707, and the edges are 32564142.

Post network construction, sparsity for all 4 networks was 99.982 % with mean absolute partial correlation of 0.0016 for the co-expression network, 0.0014 for the co-methylation network, 0.0006084186 for the multiplex network, and 0.001114587 for the monoplex network. The top 50 highly connected adjacent metrics can be found in **Figure 3** as a heatmap.

**Figure 3.**
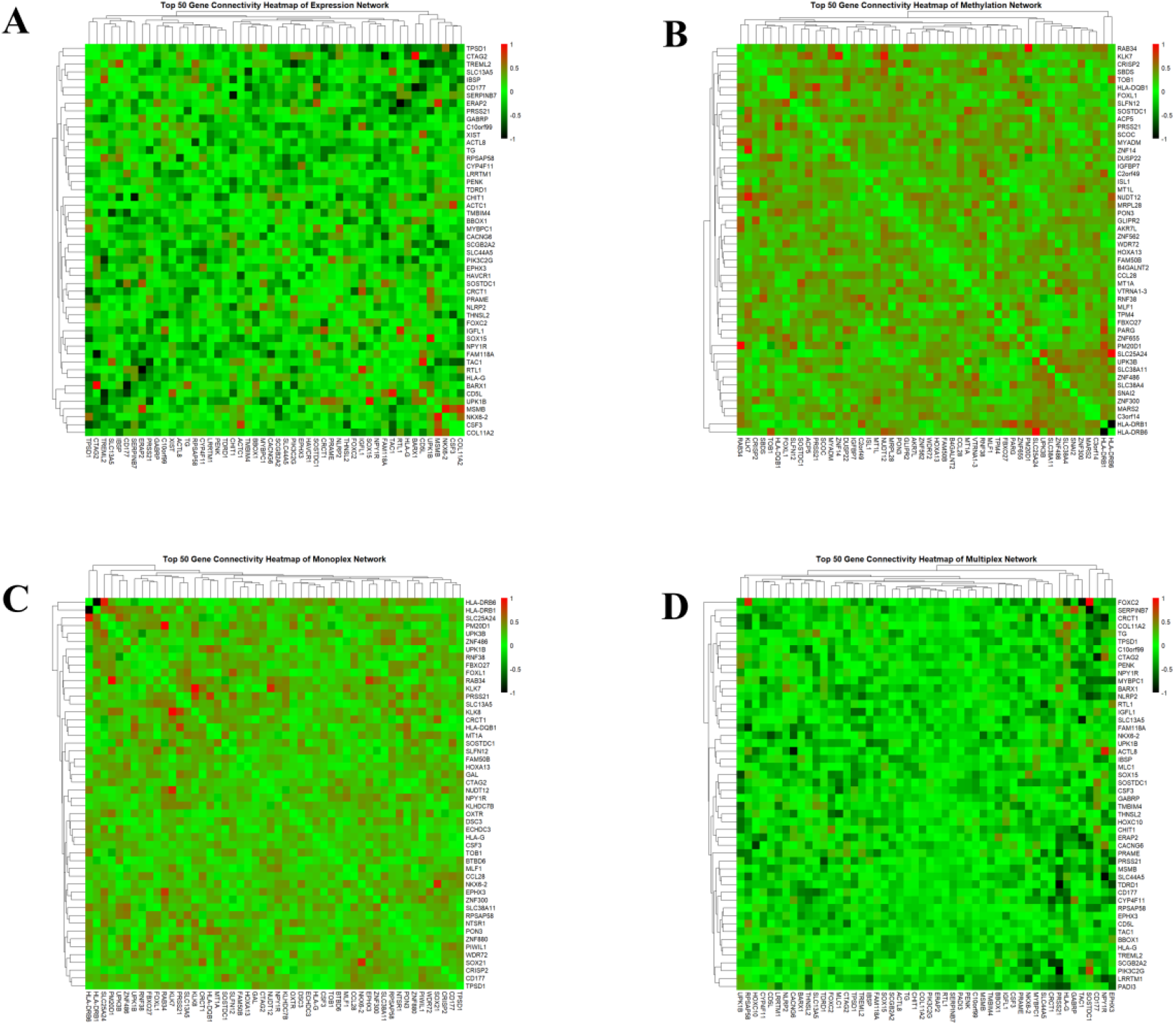
Network construction using partial correlational A) single-layer expression network, B) single-layer methylation network, C) Fused monoplex network, D) multi-layer multiplex network

### WGCNA to select the highly significant features

For the constructed partial correlation network of all 4 matrices (expression, methylation, multiplex, monoplex), WGCNA was applied to identify modules with strongly correlated genes.

Initially, a hierarchical clustering dendrogram was constructed to visualize how genes group based on their correlation patterns. As shown in **Figure 4A**, cutting this tree produced distinct clusters, and the colored bar beneath the dendrogram indicates the modules, each representing a group of highly co-expressed genes. To further correlate the relation between each module, eigengene clustering was performed to examine the relationships between modules.

**Figure 4.**
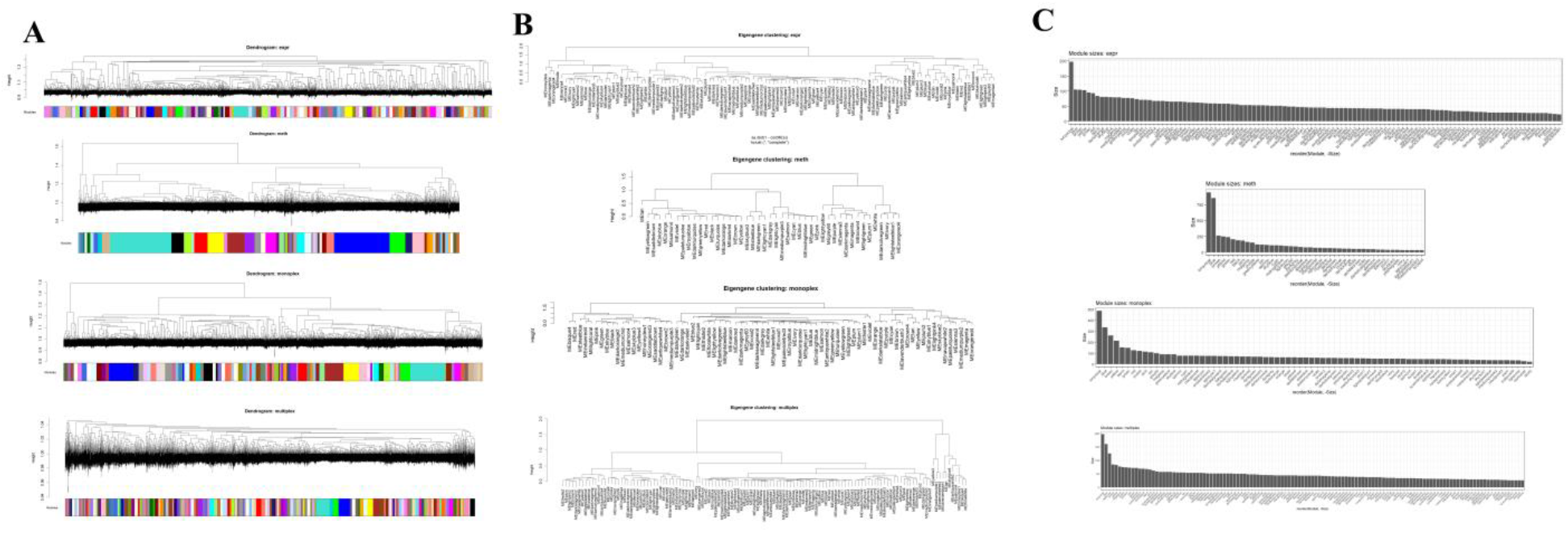
WGCNA module identification, A) Dendrogram B) Clustering of each network, C) Number of gene in each module

The expression eigengene dendrogram showed a complex and divergent network with clear branches and height. In contrast, the methylation eigengene dendrogram has a small and more compact branches, highlighting that methylation modules form tighter blocks of co-methylation with stronger internal similarity but lower overall diversity than expression modules. For the monoplex network, where expression and methylation matrices were built by fusing, show similar branch patterns of expression network. In contrast, the multiplex network keeps expression and methylation as separate layers and adds cross-layer connections, its eigengene clusters therefore summarize modules defined by both within-layer and between-layer links, highlighting groups of genes whose expression and methylation are jointly organized through these cross-layer interactions. The number of branches is more in this case compared to others, highlighting the multi-omics network storing multi-layer information. This set of four eigengene trees together illustrates how module structure changes between networks built by single omics and multi-omics data, as represented in **Figure 4B**. A detailed summary of the number of genes in each module is represented in **Figure 4C** as a bar plot.

After module clustering, eigengene correlation between each module was represented in the form of a heatmap, as illustrated in **Figure 5**. This heatmap shows how similar (correlated) the modules are to each other. orangish red colors show strong correlation, blue represents negative correlation, and yellow represents moderate correlation. Compared to single-omic networks, the multiplex integration of expression and methylation yielded substantially higher eigengene correlations across modules.

**Figure 5.**
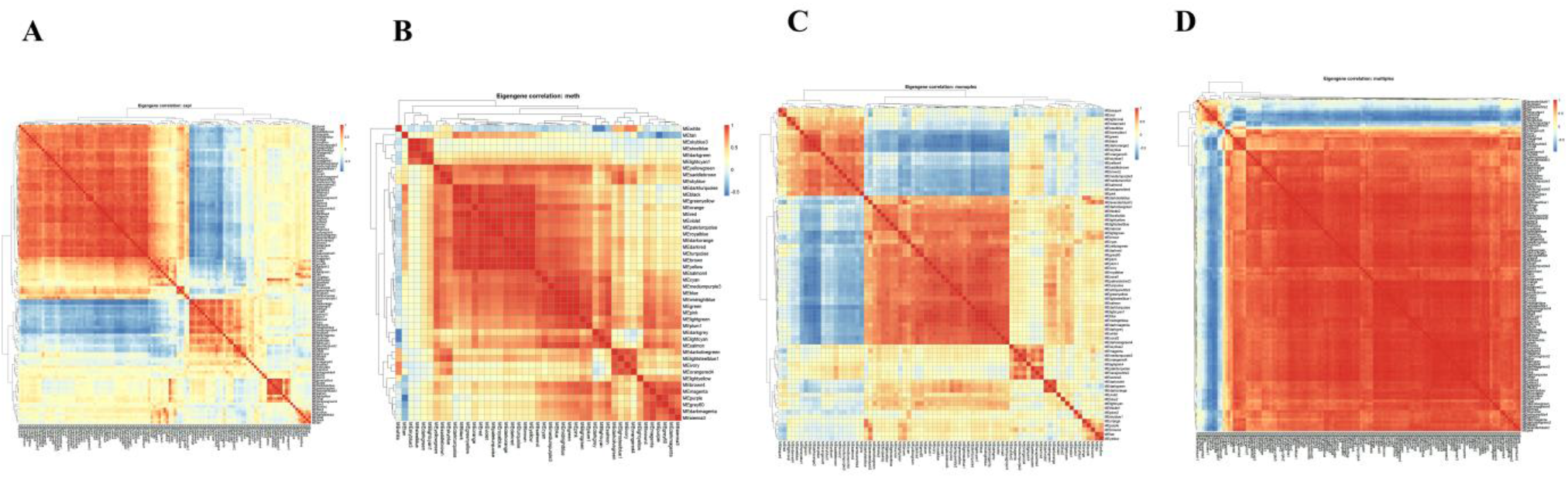
Heatmap of the top 50 gene eigengene correlation between each module

For each module, hub genes (those with high intramodular connectivity) were selected as representative features. These hub genes summarize the dominant signal of their modules and were used as inputs for machine-learning models.

### Machine learning model training and evaluation

The ROC-based evaluation of the eight datasets showed varying predictions. For models trained on the dataset of expression features with expression data, achieved AUCs between 0.58 ± 0.13 to 0.69 ± 0.09. When methylation values were added to these expression features, performance improved, with AUCs ranging from 0.68 ± 0.05 to 0.82 ± 0.07, showing that adding methylation enhances the prognostic signal in expression-based signatures. Using methylation features with added expression values resulted in an AUC of 0.54 ± 0.06 to 0.71 ± 0.11, while methylation data produced AUCs of about 0.65 ± 0.04 to 0.71 ± 0.11. The models trained on monoplex features with expression values achieved an AUC of 0.60 ± 0.08 to 0.73 ± 0.12, and monoplex methylation values reached about 0.64 to 0.77 ± 0.02, indicating that simple fusion of both omics in multi-layer modestly improves classification. The best performance was observed for multiplex-derived features: multiplex expression features produced AUCs of 0.63 ± 0.10 to 0.74 ± 0.09, whereas multiplex methylation features achieved AUCs from 0.67 ± 0.07 up to 0.82 ± 0.08. These results indicate that cross-layer connections between expression and methylation in the multiplex network capture additional information, particularly for the methylation-based multiplex features. A detailed AUC for each model for all datasets is represented in **Figure 6**.

**Figure 6.**
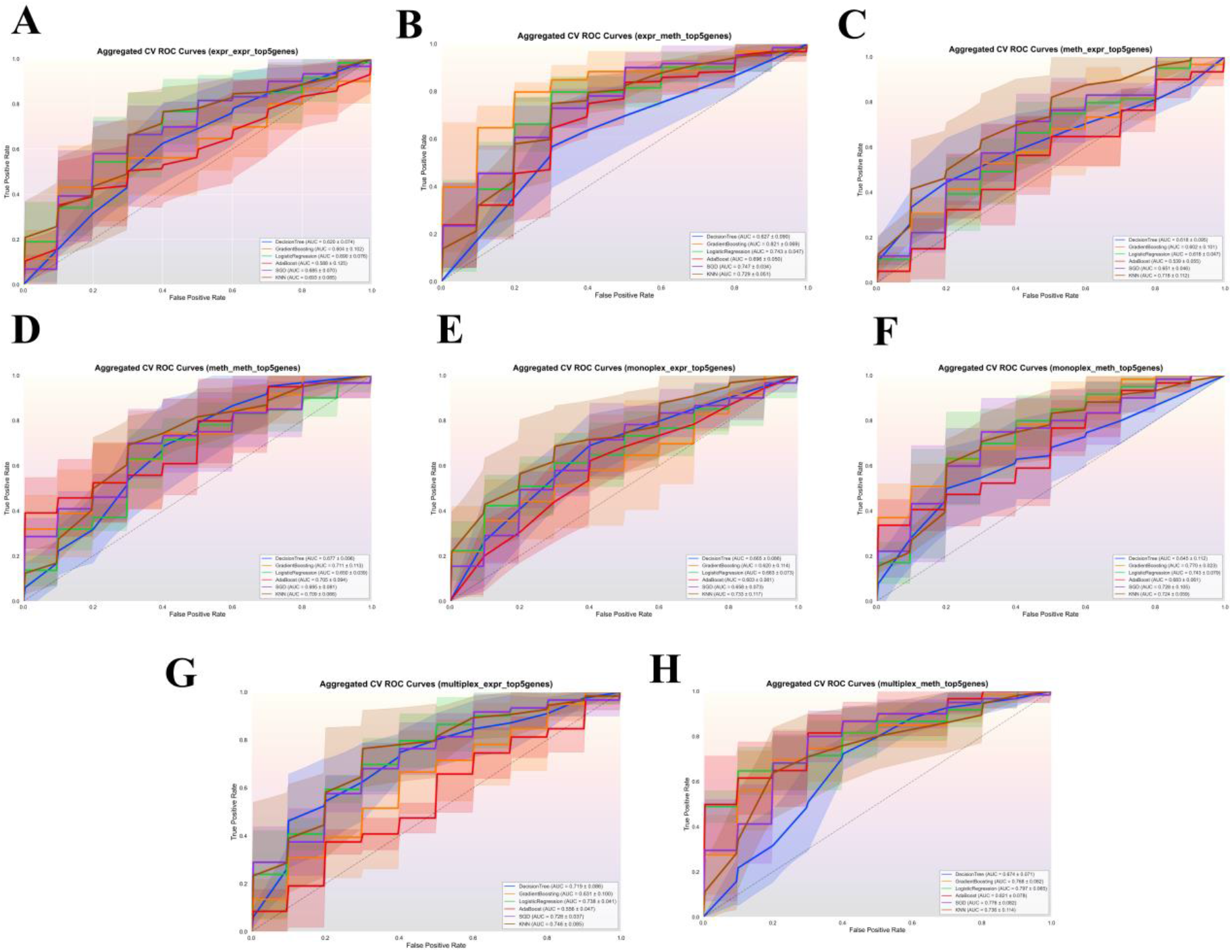
ROC curves of the eight different datasets constructed using the top 5 highly connected genes from four networks A) Classifiers trained on expression-network features using expression values, B) Classifiers trained on expression-network features using methylation values, C) Classifiers trained on methylation-network features using expression values, D) Classifiers trained on methylation-network features using methylation values, E) Classifiers trained on monoplex-network features using expression values, F) Classifiers trained on monoplex-network features using methylation values, G) Classifiers trained on multiplex-network features using expression values, H) Classifiers trained on multiplex-network features using methylation values.

Accuracy analysis showed a similar pattern to the ROC results. For models trained on expression features with expression values, accuracies ranged from 0.511 to 0.617, whereas models trained on methylation values achieved lower accuracies between 0.447 and 0.574. For methylation-derived feature sets with expression values, the accuracies improved to 0.489–0.723, indicating very good performance in several models. However, adding methylation values to these features slightly reduced accuracy to 0.574–0.660. For the monoplex representation, expression-based monoplex features yielded accuracies of 0.511–0.617, while the monoplex methylation value dataset achieved the highest accuracy, ranging from 0.553 to 0.723. Finally, for the multiplex network, the multiplex expression value dataset resulted in accuracies in the range of 0.489–0.702, whereas the multiplex methylation features produced accuracies of 0.553–0.681. Bar plots for each network are summarized in **Figure 7A**.

**Figure 7.**
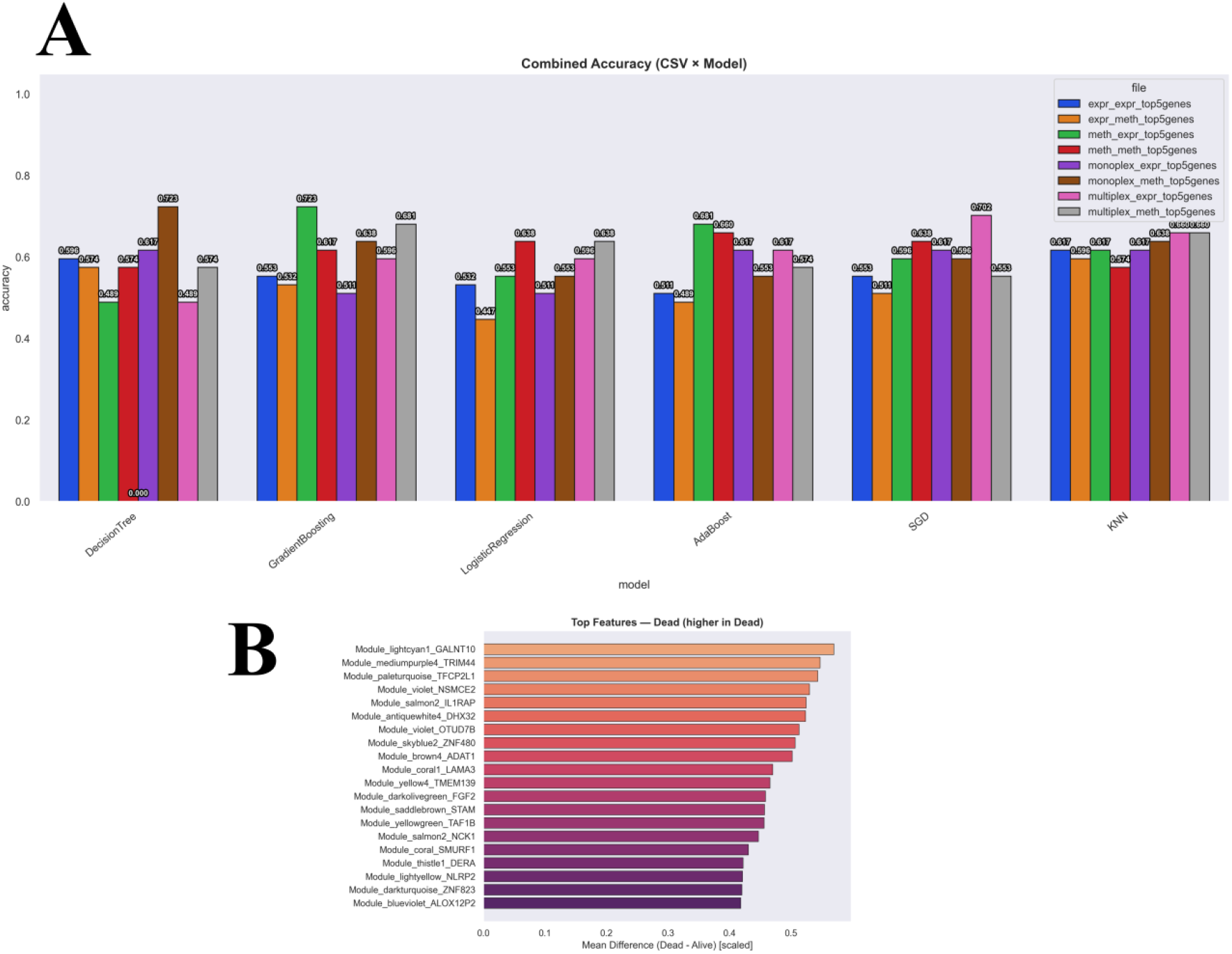
A) Accuracy plot of eight different datasets constructed using the top 5 highly connected genes from 4 networks, B) Top gene responsible for the worst survival status of the early-stage pancreatic patients

One of the important observations is that when features from one omics type are combined with values from the other, prediction accuracy increases. In contrast, when the values from the same omics were added, accuracy tended to decrease. This highlights a strong cross-omics dependency and the benefit of integrating multi-omics information rather than relying on a single data type. Taken together with the ROC and accuracy analyses, these results confirm that the best predictive performance is achieved when using multiplex features derived from both expression and methylation values. The top 20 genes contributing to the worst prognosis are shown in **Figure 7B** along with their corresponding feature importance values.

### Validation of predicted prognostic genes

To cross-validate that the predicted genes are truly associated with poor prognosis, survival analysis was performed for each gene using Kaplan-Meier (KM) plots. For each gene, the hazard ratio (HR) was calculated based on its expression patterns and corresponding clinical data, where HR > 1 indicates higher recurrence rates with poor prognosis.

For the survival analysis, patients bearing these hub genes were grouped into two groups based on the expression levels of the hub genes. A high-expression group and a low-expression group, and their survival outcomes were compared to determine which group exhibited better prognosis. Notably, all predicted genes showed HR values greater than 1, where patients with high expression of these genes had lower survival rates, confirming their association with poor prognosis. Detailed HR values and p-values for each gene are presented as a forest plot in **Figure 8A**. The Kaplan– Meier curves for the top three genes are shown in **Figure 8B–D**, highlighting a clear survival difference between the two groups. Finally, the three genes with the highest HR values, lower survival rates, and statistically significant p-values were identified as biomarkers associated with poor prognosis in early-stage pancreatic cancer: TFCP2L1, DHX32, and NCK1.

**Figure 8.**
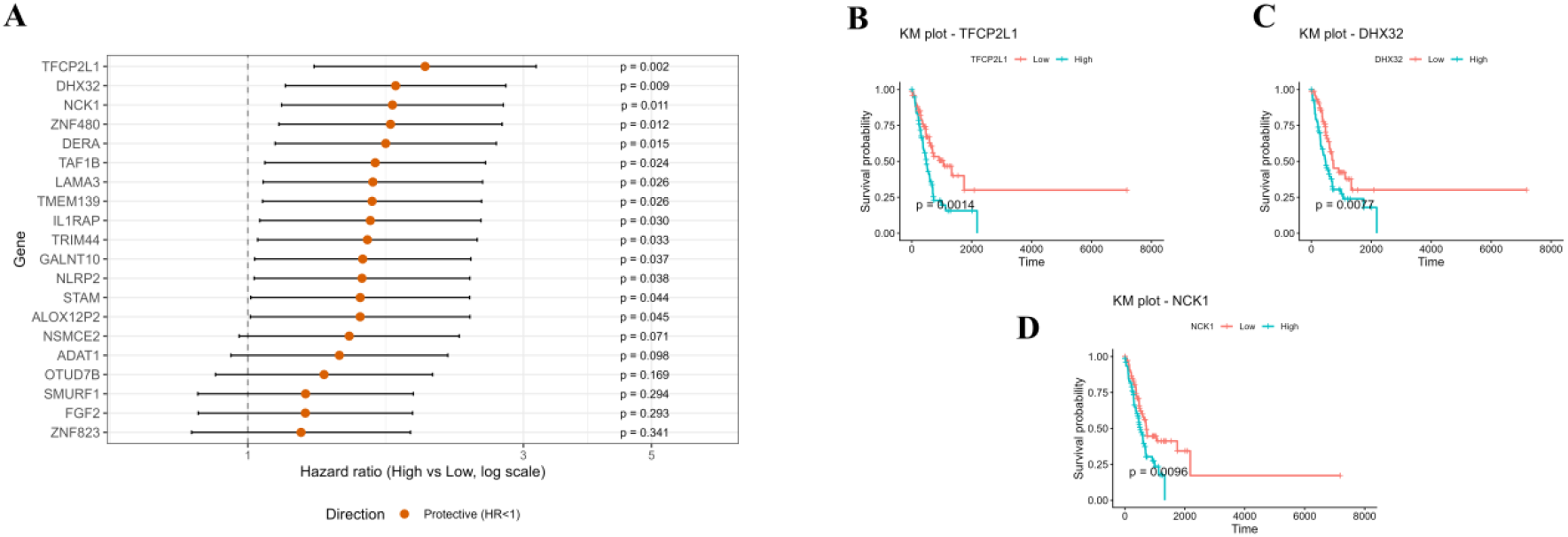
Survival analysis of top genes responsible for the poor prognosis A) HR and p value of all top 20 genes, B-D) KM plots of top hub genes.

## 4. Discussion

Early diagnosis of pancreatic cancer is often challenging due to its asymptomatic and biologically heterogeneous nature in early stages, unlike later stages, the molecular alterations are not robust or consistent. Pancreatic tumors evolve from complex epigenetic and transcriptomic changes over time, so the biomarkers identified from late-stage disease are not effective for early detection. Due to limitations, most of the present biomarkers fail in identifying the early stages pancreatic cancer accurately (Kaur et al., 2012).

Most of the studies to screen early-stage pancreatic cancer biomarkers relied on single-omics data (predominantly expression profiles), which capture only one layer of molecular information. However, pancreatic cancer is driven by the interplay of multiple omics layers between transcriptomic, epigenomic, and other regulatory layers, where biomarkers emerge from multi-entity interactions rather than isolated molecular changes (Turanli et al., 2021).

Recognizing this limitation, the present study employed a partial correlation-based multi-omics network approach that integrates both expression and methylation information to capture cross-omics interactions missed by conventional single-omics analyses, followed by hub gene prediction responsible for poor prognosis in pancreatic cancer using features derived from the constructed networks.

Before network construction, genes were ranked by variance, and genes with high variations were retained. Genes with very low variance carry little discriminative information and may induce noise into correlation networks (Parkinson et al., 2023). Therefore, they were excluded from further analysis. After intersecting high-variance genes between expression and methylation layers, 5,707 shared, highly informative genes with highly varying genes were used to construct networks.

The network is a representation of relationships between genes to show how these genes are correlated. For the construction of any network, two components are required, namely nodes and edges. Nodes represent genes, and edges represent the strength of association between genes through their values. Building these networks helps in the identification of biologically meaningful interactions (Galindez et al., 2023). Networks can be constructed using several correlation strategies, including linear (Pearson) correlation, rank-based correlations (Spearman), Kendall’s tau, and partial correlation. Here, partial correlation was used since it measures the association between two genes after accounting for the effects of all other genes in the dataset, thereby helping to distinguish direct functional relationships from indirect correlations driven by shared regulators(Masuda et al., 2025).

Using partial correlation, four types of networks were constructed. Two of them are true single-layer networks (co-expression network and the co-methylation network), where all nodes belong to a single omics layer and edges represent similarity within that layer only. A third network, the monoplex network, is also single-layer in structure because gene expression and methylation information are fused into a single adjacency matrix, where each gene is represented once, and each edge reflects an aggregated similarity combining both omics(Kosvyra et al., 2024).

However, the multiplex network is a multi-layer network, in which the same set of genes appears separately in two layers (expression and methylation). Each layer retains its own omics-specific edges, and additional cross-layer links connect corresponding genes across layers. This architecture preserves individual expression and methylation relationships while explicitly modeling the inter-layer dependency between transcriptional activity and epigenetic regulation, and these networks were used for the feature selection for ML training (Lee et al., 2020).

For training a machine-learning model, careful feature selection is important because each feature should capture a distinct and informative pattern. Adjacent matrices of the constructed four networks were used to select the features with distinct and informative information. The starting point was the network adjacency matrices, where each matrix encodes pairwise connections between genes. These adjacency matrices were first analysed using WGCNA to group genes with similar connectivity patterns into modules, and each module was treated as a distinct functional unit(Gakii et al., 2023).

However, the ultimate goal was to use individual genes rather than whole modules as features, the top five hub genes, those with the highest intramodular connectivity, were selected as representative features. This procedure was applied to all modules from each of the four networks (expression, methylation, monoplex, and multiplex). For all selected gene, expression and methylation values were kept as separate feature sets, yielding a total of eight datasets (4 networks × 2 omics). These parallel datasets enabled a systematic comparison of how module-derived genes perform when represented by expression alone, methylation alone, or their integrated network context during model training and evaluation (Gakii et al., 2023).

The machine learning results consistently demonstrated that multiplex network features provide the strongest prognostic signal compared with all other network representations. Models trained using only expression-derived features showed modest performance (AUC mostly < 0.70; accuracy 0.511–0.617), indicating that expression alone carries limited information for predicting early-stage pancreatic cancer outcomes. When methylation values were added to expression-based features, performance improved substantially (AUC 0.68–0.82; accuracy 0.574–0.660), highlighting that epigenetic information enhances the prognostic power of expression signatures.

A similar trend was observed for methylation-derived feature sets: the addition of expression features improved both AUC and accuracy (up to 0.723), confirming that the two omics layers provide complementary biological information.

Although the monoplex networks (single-layer fusion of expression and methylation) outperformed individual omics, their gains remained moderate, suggesting that simple merging of multi-omics does not fully preserve inter-layer biological structure. In contrast, multiplex network features were the most informative. Multiplex expression features surpassed the performance of single-omics and monoplex models, and multiplex methylation features achieved the best performance overall, with AUC values reaching up to 0.82 ± 0.08 and consistently strong accuracy. These findings indicate that cross-layer connectivity between expression and methylation preserves complementary biological signals that are otherwise lost in single-layer or aggregated representations.

Taken together, the results confirm that multiplex networks offer a more powerful computational framework for identifying prognostic biomarkers in early-stage pancreatic cancer than single-omics, expression-only, methylation-only, or monoplex networks. By leveraging cross-omics interactions, the multiplex approach yields more accurate survival predictions and concentrates predictive information into a focused set of genes. The top 20 genes identified through feature-importance analysis therefore represent multiplex-derived biomarkers strongly associated with poor prognosis.

Several studies have applied monoplex and multiplex network approaches for hub-gene identification. For example, in oral cancer, a multiplex network integrating expression and methylation data to predict hub genes resulted in an accuracy of about 96%, whereas hubs derived from monoplex networks achieved accuracies of approximately 88–92%. Predictions were performed with 3-nearest neighbours, Random Forest, and Support Vector Machine classifiers (Mahapatra et al., 2021).

To cross-validate that the model predictions truly reflected poor prognosis, survival analysis was performed on the top 20 candidate genes. For each gene, patients were stratified into high- and low-expression groups, with hub-gene–positive patients (high expression) forming one group and those with low expression forming the other. Kaplan–Meier survival curves were then generated to compare outcome patterns between these two groups.

Results showed that all 20 genes showed a consistent pattern in which patients with high expression had markedly worse survival than those with low expression, confirming that elevated expression of these genes is associated with poor prognosis. Finally, the three genes with the highest hazard ratios were prioritised as key hub biomarkers driving the poorest survival in early-stage pancreatic cancer. And the 3 genes include TFCP2L1, DHX32, and NCK1.

TFCP2L1 (Transcription Factor CP2 Like 1) is a transcription factor that maintains pluripotency and self-renewal in embryonic stem cells by regulating core stemness networks and cell-cycle progression (Kotarba et al., 2018). Aberrant TFCP2L1 activity has been reported in several solid tumours A broader review of the TFCP2/TFCP2L1/UBP1 family notes that TFCP2 family members, particularly TFCP2, act as pro-oncogenic factors in hepatocellular carcinoma, pancreatic cancer, and breast cancer, suggesting that TFCP2L1 may contribute to similar oncogenic transcriptional programs in pancreatic tissue (Qiu et al., 2024).

DHX32 (DEAH-box helicase 32) is an RNA helicase involved in RNA metabolism, including processes such as transcription, splicing, ribosome biogenesis, and mRNA translation. Through these functions, DHX32 influences cell proliferation, differentiation, apoptosis, and angiogenesis. DHX32 expression is dysregulated in multiple cancers: it is up-regulated in colorectal and breast cancer, where high levels promote tumour cell proliferation, migration, invasion, angiogenesis, and resistance to chemotherapy, and are associated with poor prognosis. Additionally, the Human Protein Atlas lists DHX32 as a prognostic marker in pancreatic adenocarcinoma, with altered expression associated with patient outcome, supporting its relevance in this disease (Wei et al., 2021).

NCK1 (NCK adaptor protein 1) is an SH2/SH3-containing adaptor protein that links activated receptor tyrosine kinases and other signalling proteins to downstream effectors controlling actin cytoskeleton remodelling, cell migration, proliferation, and survival. NCK1 often behaves as a pro-oncogenic adaptor (Paensuwan et al., 2022). Studies in several cancers show that NCK1 promotes proliferation, migration, invasion, and tumour angiogenesis. In pancreatic carcinoma, NCK1 has been implicated in an EGFR-dependent pathway that activates Rap1 and drives tumour cell invasion and spontaneous metastasis without affecting primary tumour growth; NCK1 knockdown blocks EGFR-induced metastasis in experimental models. Other work indicates that NCK1 integrates endoplasmic reticulum–stress signalling and can modulate cancer-cell survival, further supporting its role in tumour adaptation (Xia et al., 2019).

## Conclusion

This study demonstrates that integrating expression and methylation within a partial-correlation– based multiplex network substantially improves the identification of clinically meaningful biomarkers for early-stage pancreatic ductal adenocarcinoma. Multiplex-derived hub genes outperformed single-omics and fused monoplex features in survival prediction, indicating that explicit modelling of cross-layer dependencies captures additional prognostic signal that is otherwise missed. The prioritised genes TFCP2L1, DHX32, and NCK1 emerge as robust candidates whose network centrality and strong association with poor outcome support their potential as early prognostic markers and therapeutic targets in pancreatic cancer.

## Limitations and feature directions

The main limitations of this study are, first, the lack of external validation. All analyses were performed in a single TCGA cohort, so the multiplex hubs and performance estimates may be cohort-specific and need confirmation in independent pancreatic cancer datasets and prospective clinical series. Second, only two bulk omics layers—expression and DNA methylation—were integrated, with a relatively modest sample size for high-dimensional network and machine-learning modeling; adding further omics (such as proteomics, copy-number, or single-cell data) and larger cohorts could stabilize network topology and improve predictive accuracy. Third, the work is entirely computational, without functional experiments or assay development, so the causal roles and clinical utility of TFCP2L1, DHX32, and NCK1 remain to be demonstrated in vitro, in vivo, and in patient samples.

Future directions therefore include validating the multiplex-based signature and hub genes in multiple independent cohorts, ideally spanning different platforms and clinical settings, and extending the framework to additional omics layers to capture a more complete view of tumour biology. Experimental follow-up should test the mechanistic contribution of TFCP2L1, DHX32, and NCK1 to pancreatic cancer progression and treatment response, and support development of practical assays—such as ELISA or multiplex immunoassays—for measuring these markers in blood or tissue for use in early prognostic stratification.

## References

1. Ballehaninna, U. K., & Chamberlain, R. S. (2012). The clinical utility of serum CA 19-9 in the diagnosis, prognosis and management of pancreatic adenocarcinoma: An evidence based appraisal. Journal of Gastrointestinal Oncology, 3(2), 105. 10.3978/J.ISSN.2078-6891.2011.021

2. Barkeer, S., Chugh, S., Batra, S. K., & Ponnusamy, M. P. (2018). Glycosylation of Cancer Stem Cells: Function in Stemness, Tumorigenesis, and Metastasis. Neoplasia, 20(8), 813–825. 10.1016/J.NEO.2018.06.001

3. Black, D., Byrne, D., Walke, A., Liu, S., Di Ieva, A., Kaneko, S., Stummer, W., Salcudean, T., & Suero Molina, E. (2024). Towards machine learning-based quantitative hyperspectral image guidance for brain tumor resection. Communications Medicine, 4(1), 131. 10.1038/S43856-024-00562-3

4. Boldini, D., Grisoni, F., Kuhn, D., Friedrich, L., & Sieber, S. A. (2023). Practical guidelines for the use of gradient boosting for molecular property prediction. Journal of Cheminformatics, 15(1), 73. 10.1186/S13321-023-00743-7

5. Chai, K., Liang, J., Zhang, X., Cao, P., Chen, S., Gu, H., Ye, W., Liu, R., Hu, W., Peng, C., Liu, G. L., & Shen, D. (2021). Application of Machine Learning and Weighted Gene Co-expression Network Algorithm to Explore the Hub Genes in the Aging Brain. Frontiers in Aging Neuroscience, 13, 707165. 10.3389/FNAGI.2021.707165/FULL

6. Cheadle, C., Vawter, M. P., Freed, W. J., & Becker, K. G. (2003). Analysis of Microarray Data Using Z Score Transformation. The Journal of Molecular Diagnostics : JMD, 5(2), 73. 10.1016/S1525-1578(10)60455-2

7. Fraunhoffer, N. A., Abuelafia, A. M., Bigonnet, M., Gayet, O., Roques, J., Nicolle, R., Lomberk, G., Urrutia, R., Dusetti, N., & Iovanna, J. (2022). Multi-omics data integration and modeling unravels new mechanisms for pancreatic cancer and improves prognostic prediction. Npj Precision Oncology 2022 6:1, 6(1), 57-. 10.1038/s41698-022-00299-z

8. Gakii, C., Mukami, V., & Too, B. (2023). Feature selection for classification using WGCNA and Spread Sub-Sample for an imbalanced rheumatoid arthritis RNASEQ data. Informatics in Medicine Unlocked, 43, 101402. 10.1016/J.IMU.2023.101402

9. Galindez, G., Sadegh, S., Baumbach, J., Kacprowski, T., & List, M. (2023). Network-based approaches for modeling disease regulation and progression. Computational and Structural Biotechnology Journal, 21, 780–795. 10.1016/J.CSBJ.2022.12.022

10. Gomes Mantovani, R., Horváth, T., Rossi, A. L. D., Cerri, R., Barbon Junior, S., Vanschoren, J., & Carvalho, A.C.P.L.F.de. (2024). Better trees: an empirical study on hyperparameter tuning of classification decision tree induction algorithms. Data Mining and Knowledge Discovery 2024 38:3, 38(3), 1364–1416. 10.1007/S10618-024-01002-5

11. Jensen, M. A., Ferretti, V., Grossman, R. L., & Staudt, L. M. (2017). The NCI Genomic Data Commons as an engine for precision medicine. Blood, 130(4), 453. 10.1182/BLOOD-2017-03-735654

12. Jiang, W., Ye, W., Tan, X., & Bao, Y. J. (2025). Network-based multi-omics integrative analysis methods in drug discovery: a systematic review. BioData Mining, 18(1), 27. 10.1186/S13040-025-00442-Z

13. Kaur, S., Baine, M. J., Jain, M., Sasson, A. R., & Batra, S. K. (2012). Early diagnosis of pancreatic cancer: challenges and new developments. Biomarkers in Medicine, 6(5), 597. 10.2217/BMM.12.69

14. Kim, H. S., Choi, Y. H., Lee, J. S., Jo, I. H., Ko, S. W., Paik, K. H., Choi, H. H., Lee, H. H., Lim, Y. S., Paik, C. N., Lee, I. S., & Chang, J. H. (2024). Characteristics of Early Pancreatic Cancer: Comparison between Stage 1A and Stage 1B Pancreatic Cancer in Multicenter Clinical Data Warehouse Study. Cancers, 16(5), 944. 10.3390/CANCERS16050944

15. Ko, D. K., & Brandizzi, F. (2020). Network-based approaches for understanding gene regulation and function in plants. The Plant Journal : For Cell and Molecular Biology, 104(2), 302. 10.1111/TPJ.14940

16. Kosvyra, Karadimitris, Papaioannou, & Chouvarda, I. (2024). Machine learning and integrative multi-omics network analysis for survival prediction in acute myeloid leukemia. Computers in Biology and Medicine, 178, 108735. 10.1016/J.COMPBIOMED.2024.108735

17. Kotarba, G., Krzywinska, E., Grabowska, A. I., Taracha, A., & Wilanowski, T. (2018). TFCP2/TFCP2L1/UBP1 transcription factors in cancer. Cancer Letters, 420, 72–79. 10.1016/J.CANLET.2018.01.078

18. Kuijpers, T. J. M., Kleinjans, J. C. S., & Jennen, D. G. J. (2021). From multi-omics integration towards novel genomic interaction networks to identify key cancer cell line characteristics. Scientific Reports 2021 11:1, 11(1), 10542-. 10.1038/s41598-021-90047-3

19. Langfelder, P., & Horvath, S. (2007). Eigengene networks for studying the relationships between co-expression modules. BMC Systems Biology, 1, 54. 10.1186/1752-0509-1-54

20. Lee, B., Zhang, S., Poleksic, A., & Xie, L. (2020). Heterogeneous Multi-Layered Network Model for Omics Data Integration and Analysis. Frontiers in Genetics, 10, 501269. 10.3389/FGENE.2019.01381/BIBTEX

21. Leiphrakpam, P. D., Chowdhury, S., Zhang, M., Bajaj, V., Dhir, M., & Are, C. (2025). Trends in the Global Incidence of Pancreatic Cancer and a Brief Review of its Histologic and Molecular Subtypes. Journal of Gastrointestinal Cancer, 56(1). 10.1007/S12029-025-01183-2

22. Liang, J., & Tong, W. G. (2023). Ultrasensitive Detection and Separation of Pancreatic Cancer Biomarker CA 19-9 Using a Multiphoton Laser Wave-Mixing Detector Interfaced to Capillary Electrophoresis. ACS Omega, 8(34), 31030–31039. 10.1021/ACSOMEGA.3C02845

23. Mahapatra, S., Bhuyan, R., Das, J., & Swarnkar, T. (2021). Integrated multiplex network based approach for hub gene identification in oral cancer. Heliyon, 7(7), e07418. 10.1016/J.HELIYON.2021.E07418

24. Masuda, N., Boyd, Z. M., Garlaschelli, D., & Mucha, P. J. (2025). Introduction to correlation networks: Interdisciplinary approaches beyond thresholding. Physics Reports, 1136, 1–39. 10.1016/J.PHYSREP.2025.06.002

25. Paensuwan, P., Ngoenkam, J., Wangteeraprasert, A., & Pongcharoen, S. (2022). Essential function of adaptor protein Nck1 in platelet-derived growth factor receptor signaling in human lens epithelial cells. Scientific Reports 2022 12:1, 12(1), 1063-. 10.1038/s41598-022-05183-1

26. Pan, C., Tang, H., Wang, W., Wu, D., Luo, H., Xu, L., & Lin, X. J. (2023). An enhanced genetic mutation-based model for predicting the efficacy of immune checkpoint inhibitors in patients with melanoma. Frontiers in Oncology, 12. 10.3389/FONC.2022.1077477

27. Parkinson, E., Liberatore, F., Watkins, W. J., Andrews, R., Edkins, S., Hibbert, J., Strunk, T., Currie, A., & Ghazal, P. (2023). Gene filtering strategies for machine learning guided biomarker discovery using neonatal sepsis RNA-seq data. Frontiers in Genetics, 14, 1158352. 10.3389/FGENE.2023.1158352/FULL

28. Qiu, D., Wang, T., Xiong, Y., Li, K., Qiu, X., Feng, Y., Lian, Q., Qin, Y., Liu, K., Zhang, Q., & Jia, C. (2024). TFCP2L1 drives stemness and enhances their resistance to Sorafenib treatment by modulating the NANOG/STAT3 pathway in hepatocellular carcinoma. Oncogenesis, 13(1). 10.1038/S41389-024-00534-1

29. Siddalingappa, R., & Kanagaraj, S. (2023). K-nearest-neighbor algorithm to predict the survival time and classification of various stages of oral cancer: a machine learning approach. F1000Research, 11, 70. 10.12688/F1000RESEARCH.75469.2

30. Singhi, A. D., Koay, E. J., Chari, S. T., & Maitra, A. (2019). Early Detection of Pancreatic Cancer: Opportunities and Challenges. Gastroenterology, 156(7), 2024. 10.1053/J.GASTRO.2019.01.259

31. Turanli, B., Yildirim, E., Gulfidan, G., Arga, K. Y., & Sinha, R. (2021). Current State of “Omics” Biomarkers in Pancreatic Cancer. Journal of Personalized Medicine, 11(2), 127. 10.3390/JPM11020127

32. Wei, Q., Geng, J., Chen, Y., Lin, H., Wang, J., Fang, Z., Wang, F., & Zhang, Z. (2021). Structure and function of DEAH-box helicase 32 and its role in cancer. Oncology Letters, 21(5), 382. 10.3892/OL.2021.12643

33. Xia, P., Huang, M., Zhang, Y., Xiong, X., Yan, M., Xiong, X., Yu, W., & Song, E. (2019). NCK1 promotes the angiogenesis of cervical squamous carcinoma via Rac1/PAK1/MMP2 signal pathway. Gynecologic Oncology, 152(2), 387–395. 10.1016/J.YGYNO.2018.11.013

34. Xu, W., Xu, M., Wang, L., Zhou, W., Xiang, R., Shi, Y., Zhang, Y., & Piao, Y. (2019). Integrative analysis of DNA methylation and gene expression identified cervical cancer-specific diagnostic biomarkers. Signal Transduction and Targeted Therapy 2019 4:1, 4(1), 55-. 10.1038/s41392-019-0081-6

35. Yang, C. Y., Lin, R. T., Chen, C. Y., Yeh, C. C., Tseng, C. M., Huang, W. H., Lee, T. Y., Chu, C. S., & Lin, J. T. (2022). Accuracy of simultaneous measurement of serum biomarkers: Carbohydrate antigen 19-9, pancreatic elastase-1, amylase, and lipase for diagnosing pancreatic ductal adenocarcinoma. Journal of the Formosan Medical Association, 121(12), 2601–2607. 10.1016/J.JFMA.2022.07.003

36. Zhang, Z., Trevino, V., Hoseini, S. S., Belciug, S., Boopathi, A. M., Zhang, P., Gorunescu, F., Subha, V., & Dai, S. (2018). Variable selection in Logistic regression model with genetic algorithm. Annals of Translational Medicine, 6(3), 45. 10.21037/ATM.2018.01.15

